# G-CSF Secreted by Epigenetically Reprogrammed Mutant IDH1 Glioma Stem Cells Reverses the Myeloid Cells’-Mediated Immunosuppressive Tumor Microenvironment

**DOI:** 10.1101/2020.07.22.215954

**Authors:** Mahmoud S Alghamri, Ruthvik P Avvari, Rohit Thalla, Neha Kamran, Li Zhang, Maria Ventosa, Ayman Taher, Syed Mohd Faisal, Felipe J. Núñez, María Belén Garcia-Fabiani, Santiago Haase, Stephen Carney, Daniel Orringer, Shawn Hervey-Jumper, Jason Heth, Parag G Patil, Wajd N Al-Holou, Karen Eddy, Sophia Merajver, Peter J Ulintz, Joshua Welch, Chao Gao, Jialin Liu, Gabriel Núñez, Dolores Hambardzumyan, Pedro R Lowenstein, Maria G Castro

## Abstract

Mutation in isocitrate dehydrogenase (*mIDH*) is a gain of function mutation resulting in the production of the oncometabolite, R-2-hydroxyglutarate, that inhibits DNA and histone demethylases. The resultant hypermethylation phenotype reprograms the glioma cells’ transcriptome and elicits profound effects on glioma immunity. We report that in mouse models and human gliomas, *mIDH1* in the context of *ATRX* and *TP53* inactivation results in global expansion of the granulocytic myeloid cells’ compartment. Single-cell RNA-sequencing coupled with mass cytometry analysis revealed that these granulocytes are mainly non-immunosuppressive neutrophils and pre-neutrophils; with a small fraction of polymorphonuclear myeloid-derived suppressor cells. The mechanism of *mIDH1* mediated pre-neutrophils expansion involves epigenetic reprogramming which leads to enhanced expression of the granulocyte colony-stimulating factor (G-CSF). Blocking G-CSF restored the inhibitory potential of PMN-MDSCs and enhanced tumor progression. Thus, G-CSF induces remodeling of the inhibitory PMN-MDSCs in *mIDH1* glioma rendering them non-immunosuppressive; and having significant therapeutic implications.

**SIGNIFICANCE:** *mIDH1* is the most common mutation in gliomas associated with improved prognosis. Gliomas harboring *mIDH1*, together with *ATRX* and *TP53* inactivation, exhibit higher circulating levels of G-CSF, ensuing the recruitment and expansion of non-suppressive neutrophils, pre-neutrophils and small fraction of PMN-MDSCs to the TME leading to an immune permissive phenotype.

## INTRODUCTION

Mutation in isocitrate dehydrogenase (*mIDH*) is a common genetic lesion occurring in adult glioma patients (1). Approximately 90% of *IDH1* mutations occur in exon 4 at codon 132, resulting in a change of a single amino acid from arginine to histidine (R132H). Less common *IDH2* mutations occur in an analogous codon at position R172 (2). Although IDH1/2 mutations are always heterozygous, they exert a dominant gain of function enzymatic activity which leads to the production of 2-hydroxyglutarate (2HG). Excessive 2HG production causes DNA hypermethylation via inhibition of methylcytosine dioxygenase *TET2* (3,4), and also promotes histone hypermethylation through competitive inhibition of α-ketoglutarate (αKG)-dependent Jumonji-C histone demethylases (5,6). This leads to epigenetic reprogramming of gene expression within the *mIDH1* glioma cells (5,7,8).

The *IDH1* mutation plays an important role in glioma development and progression, by modulating tumor cells’ intrinsic mechanisms and also by remodeling the tumor microenvironment. Metabolically, IDH1 is one of the enzymes that encodes an irreversible reaction in the tricyclic acid cycle. Disruption of the IDH reaction results in defective mitochondrial oxidative phosphorylation, glutamine metabolism, lipogenesis, glucose sensing, and altering cellular redox status (9–14). Mutation in *IDH1* also inhibits glioma stem cell differentiation (5,13), upregulates vascular endothelial growth factor (15,16), and produces high levels of hypoxia-inducible factor-1α, all of which promote glioma invasion (14,17,18). We have recently shown that *mIDH1* when expressed concomitantly with *ATRX* and *TP53* inactivation, results in enhanced DNA damage response, leading to radio-resistance in mouse and human glioma cells *in vitro* and extending the median survival (MS) in a genetically engineered mouse glioma model (19). Nevertheless, the molecular mechanisms that mediate enhanced survival in patients with *mIDH1* glioma remain elusive. We hypothesize that epigenetic-mediated mechanisms could be an important contributing factor modulating the immune microenvironment

The success of immunotherapeutic approaches in several non-central nervous system (CNS) cancers has driven the evaluation of numerous immune-mediated therapeutic strategies in glioma clinical trials (20). To date, Immune checkpoint blockade used as a monotherapy in combination with standard of care did not demonstrate survival benefits in glioma patients (21–23). The lack of therapeutic efficacy of immunotherapies in the clinical arena, can be attributed in part, to the immunosuppressive properties elicited by glioma infiltrating immune cells (22–24). Myeloid-derived suppressor cells (MDSCs) have emerged as one of the dominant immunosuppressive cells that directly interfere with the efficacy of immunotherapy (25–27). In glioma patients, it has been demonstrated that granulocytic-MDSCs (also known as PMN-MDSCs) are a major subset that expands during glioma progression which negatively correlates with patient’s survival (28–30). We have recently shown that depletion of MDSCs in wild-type *IDH1* glioma-bearing mice markedly enhanced the efficacy of an immune stimulatory/conditional cytotoxic gene therapy (25); highlighting the role of MDSCs in hampering anti-glioma immunity.

Several studies suggested that *mIDH1* may play a critical role in shaping the immunological landscape of the tumor microenvironment (31–34). Glioma samples from patients with *mIDH1* have decreased PD-L1 expression (due to hypermethylation of the CD274 promoter) (32), reduced level of inflammation, and reduced levels of infiltrating immune cells (31,33,35). Myeloid cells represent the major immune cells in glioma TME (25,36,37). It has been shown that *mIDH1* glioma with mixed genetic background (*PDGFB/shP53/ Ink4a/Arf^-/-^/mIDH1*) exhibit fewer tumor-infiltrating macrophages, dendritic cells (DCs), and neutrophils, compared to the wild-type *IDH1* counterparts (31). Nevertheless, the impact of *mIDH1* on tumor-infiltrating myeloid cells’ phenotype and function have not been explored; particularly in the context of concurrent inactivation mutation in *ATRX* and *TP53* (1,38).

Granulocytes are the major population within the bone marrow (BM) myeloid cells’ compartment, which differentiate under steady-state conditions into neutrophils. Under physiological conditions, these cells have emerged as major contributors to protection against pathogens and are important mediators of tissue remodeling (39,40). However, during inflammation and cancer, the activation of various signaling pathways, such as Jak-Stat, and MAPK pathways, results in rapid mobilization of granulocytes and neutrophils from the BM, favoring the generation of immunosuppressive PMN-MDSCs (41–43). These cells share the same origin (granulocyte-macrophage progenitors (GMPs)) and differentiation pathways, yet, PMN-MDSCs have distinct genomic and biochemical features and are immunosuppressive (44,45). Therefore, it became critical to incorporate both transcriptomic analysis at the single-cell level (single-cell RNA sequencing) as well as functional characterization to uncover the immunosuppressive tumor-infiltrating PMN-MDSCs in glioma.

We report that in genetically engineered mouse glioma models, *IDH1* mutation, in the context of *ATRX* and *TP53* deficiency, caused an expansion of tumor-infiltrating granulocytes.

Upon phenotypic and functional characterization, we uncovered that granulocytes in *mIDH1* glioma, counter to the granulocytes infiltrated in *wtIDH1*, did not exhibit immune-suppressive properties. Single-cell sequencing coupled with mass cytometry analysis revealed that these granulocytes are heterogeneous and composed of neutrophils and pre-neutrophils, with a smaller proportion of *bona fide* immunosuppressive PMN-MDSCs. Moreover, primary human gliomas showed a higher frequency of cells exhibiting the PMN-MDSCs gene signature in *wtIDH1* tumors when compared to *mIDH1* glioma. Our data demonstrate that the mechanism by which *mIDH1* in glioma mediates the expansion of non-immune suppressive granulocytes involves epigenetic reprogramming. This leads to enhanced expression of granulocyte colony-stimulating factor (G-CSF) by stem-like *mIDH1* glioma cells. Blocking G-CSF restored the inhibitory potential of PMN-MDSCs and substantially enhanced tumor progression, whereas recombinant G-CSF administration, prolonged the MS of *wtIDH1* glioma bearing mice. Consistent with that, LGG-astrocytoma with *mIDH1* is the sole tumor cohort within all TCGA data sets in which high *CSF3* gene expression correlates with favorable patients’ outcome. Our work provides a new insight of the impact of *mIDH1* on the phenotypic and functional properties of myeloid cells in the glioma TME.

## MATERIALS AND METHODS

### Study design

To study the impact of glioma *IDH1^R123H^* on myeloid cell phenotype and function, we generated a genetically engineered animal model injecting SB plasmids encoding NRA G12V, *shATRX* and *shp53* with or without *IDH1^R132H^* (*wtIDH1* and *mIDH1* respectively) into the lateral ventricle of neonatal mice (46–48). Sample size and any data inclusion/exclusion were defined individually for each experiment. We also used an animal model generated by intracranial implantation of glioma NS (*wtIDH1* and *mIDH1*) derived from our genetically engineered animal model (46,49). Additionally, we used an alternative model (PDGFB/shP53/shATRX/Ink4a/Arf^-/-^ *wtIDH1*-NS or *mIDH1*-NS) that does not encode RAS-activating mutations. We performed pan characterization of all immune cells by flow cytometry and mass cytometry and identified each immune cell population using unique biomarkers. Detailed information regarding the source of antibodies and immune cells’ gating are listed in supplementary tables S1, S2, and S3. We also performed scRNA-seq to define the PMN-MDSCs cell population in both mouse and human glioma tissue derived from patients harboring *IDH1^R132H^* or *wtIDH1*. We validated PMN-MDSCs by functional analysis of T-cell’s inhibitory characteristics. The numbers of replicates are reported in the figure legends. All scRNA-seq data were deposited in public databases as is indicated in the respective sections. Materials and Methods are described in detail in the Supplementary Materials section.

### Statistical analysis

Sample sizes were selected based on preliminary data from pilot experiments and previously published results in the literature and our laboratory. Unpaired Student t-test or one-way analysis of variance (ANOVA), followed by Tukey’s multiple comparisons post-test were utilized for comparing experimental groups with controls from flow cytometry analysis, cytokine analysis, mass cytometry analysis, and T-cell functional assays. RNA-seq, data were processed using the Tuxedo Suite, and differentially expressed genes were considered when FDR ≤ 0.05, and fold change ≥ ± 1.5. All single-cell sequencing data were processed by Cell Ranger Pipeline version 3.1.0. Filtered digital gene expression matrix files (DGE) matrices containing the number of unique molecular identifiers (UMI) counts per gene per cell were analyzed using the Seurat R package version 3.1.4. Statistically significant principal components were determined using the jackstraw function (50,51). All quantitative data are presented as the Means ± SEM from at least three independent samples. ANOVA and two-sample t-tests were used to compare continuous outcomes between groups. Survival curves were analyzed using the Kaplan-Meier method and compared using Mantel-Cox tests; the effect size is expressed as MS. Differences were considered significant if *P*< 0.05. All analyses were conducted using GraphPad Prism software (version 8.00), or R (version 3.4). The statistical tests used are indicated in each figure legend.

## RESULTS

### *mIDH1* glioma causes a systemic increase in polymorphonuclear (PMN) myeloid cells

The role of *IDH1* mutation in shaping the immune landscape of glioma remains unexplained. We generated a *mIDH1* genetically engineered mouse model (GEMM) of glioma using the Sleeping Beauty (SB) transposon system (46,52,53) to uncover the impact of *IDH1^R132H^* in the context of ATRX and TP53 deficiency on glioma infiltrating immune cells (Fig. 1A). Two groups of tumors were generated using the SB method, each contains a combination of genetic lesions commonly encountered in astrocytoma; *mIDH1* (*NRASG12V*+ *shp53* + *shATRX* + *mIDH1^R132H^*) and control *wtIDH1* group (*NRASG12V*+ *shp53* + *shATRX* + *wtIDH1*) (Fig. 1B). We used the merit of mass cytometry (CyTOF) (Supplementary Table S1) which measures 37 parameters simultaneously to analyze immune cell composition within the TME from each group. Using SPADE analysis, we found that glioma TME contains several immune cell types including macrophages, dendritic cells (DCs), B-cells, NK cells, CD8^+^ T-cells, and CD4^+^ T-cells (Fig. 1C, D). Moreover, there was an expansion of cells (depicted by the large red/yellow nodes) characteristic for myeloid-derived suppressor cells (CD45^high^/CD11b^+^/Gr-1^+^) in the TME of *mIDH1* glioma compared to *wtIDH1* glioma (Fig. 1C, D). Similarly, visual inspection of viSNE plots, which represent high-dimensional CyTOF output in two dimensions (54), showed a higher percentage of Gr-1^+^ cells in the TME from *mIDH1* glioma compared to those from *wtIDH1* glioma (Fig. 1C, D; insets). We confirmed that this difference is statistically significant using a *χ*^2^ test (*p* < 0.0001). The Gr-1 antibody can detect both the granulocytic (CD45^high^/CD11b^+^/Ly6G^+^/Ly6C^low^) and monocytic (CD45^high^/CD11b^+^/Ly6G^-^/Ly6C^high^) MDSCs population. We found that the majority of the expanded myeloid cells in *mIDH1* tumor belonged to the granulocytic population, with no changes in the frequency of the monocytic population (Supplementary Fig S1A). In addition to the increase in granulocytes, *mIDH1* tumors showed a decreased frequency of macrophages (2.7 fold) and dendritic cells ((DCs) ~ 4 fold) with no significant change in the frequency of lymphocytes (including NK cells, B-cells, CD8^+^ and CD4^+^ T-cells) as compared to *wtIDH1* tumors (Supplementary Fig. S1B-G). Notably, there was no significant difference in the percentage of CD45^+^ cells between *wtIDH1* and *mIDH1* glioma (Supplementary Fig. S1H). We generated neurospheres (NS) from mouse glioma subgroups (Fig 1B), and we used the implantable animal model to analyze the frequency of CD45^high^/CD11b^+^/Gr-1^+^ population at mid-stage (12 days after implantation) and at symptomatic stage of tumor development (Fig 1E, F). In both time points, we found that the *mIDH1* tumor group had an increased number and percentage of CD45^high^/CD11b^+^/Gr-1^+^ cells compared to the *wtIDH1* group (Mid-stage: 6.91 ± 1.4% vs. 17.52 ± 3.6%, Symptomatic stage: 32.5 ± 4.07% vs. 73.9 ± 4.6% in *wtIDH1*vs *mIDH1*; respectively, Fig. 1E, F; Supplementary Fig. S2A); the majority of these cells belong to granulocytic (Ly6G^+^) origin (Fig. 1G). We validated our results using an animal model of *mIDH1* glioma independent of *RAS* activating mutation (36,55). Brain tumors were induced with replication-competent avian leukemia virus splice acceptor (RCAS) expressing platelet-derived growth factor β (*PDGFB*), *shP53*, and *mIDH1* or *wtIDH1* in mixed background *NTva*_*Ink4a*/*Arf* ^-/-^ mice. The neurospheres generated from these tumors were engineered to encode *shATRX* to generate glioma cells with the following molecular alterations: *PDGFB/shP53/shATRX/Ink4a/Arf^-/-^/mIDH1* or *PDGFB/shP53/shATRX/Ink4a/Arf^-/-^/wtIDH1* (46). Using this additional GEMM, we confirmed that *mIDH1* tumor-bearing animals exhibit an expanded CD45^high^/CD11b^+^/Gr-1^+^ population with a high predominance of CD45^high^/CD11b^+^/Ly6G^+^ population (~2-fold, *P*<0.001; Supplementary Fig. S2D, E). The expansion of CD45^high^/CD11b^+^/Gr-1^+^ cells was not limited to the tumor site; in both the circulation and the spleen of *mIDH1* tumor-bearing animals, there was a higher (*P<0.005*) number and percentage of CD45^high^/CD11b^+^/Gr-1^+^ (Blood; 17.65 ± 2.3% vs. 50.05 ± 4.4%, spleen; 26.6 ± 2.7 vs. 66.8 ± 5.4% in *wtIDH1* vs *mIDH1* tumor, respectively), the majority of which belong to granulocytic origin (Fig. 1H-M, Supplementary Fig. S2B, C). Similar to the TME, in the circulation and spleen from tumor-bearing animals, there was no significant difference in the M-MDSCs between *wtIDH1* and *mIDH1* glioma bearing mice. However, *mIDH1* exhibited a decreased number of macrophages, with no differences in the DCs or lymphocytes (Supplementary Fig. S3, 4). Collectively, these results suggest that *mIDH1* tumor causes activation of pathways involved in granulocytic myeloid cells’ expansion.

**Figure 1.**
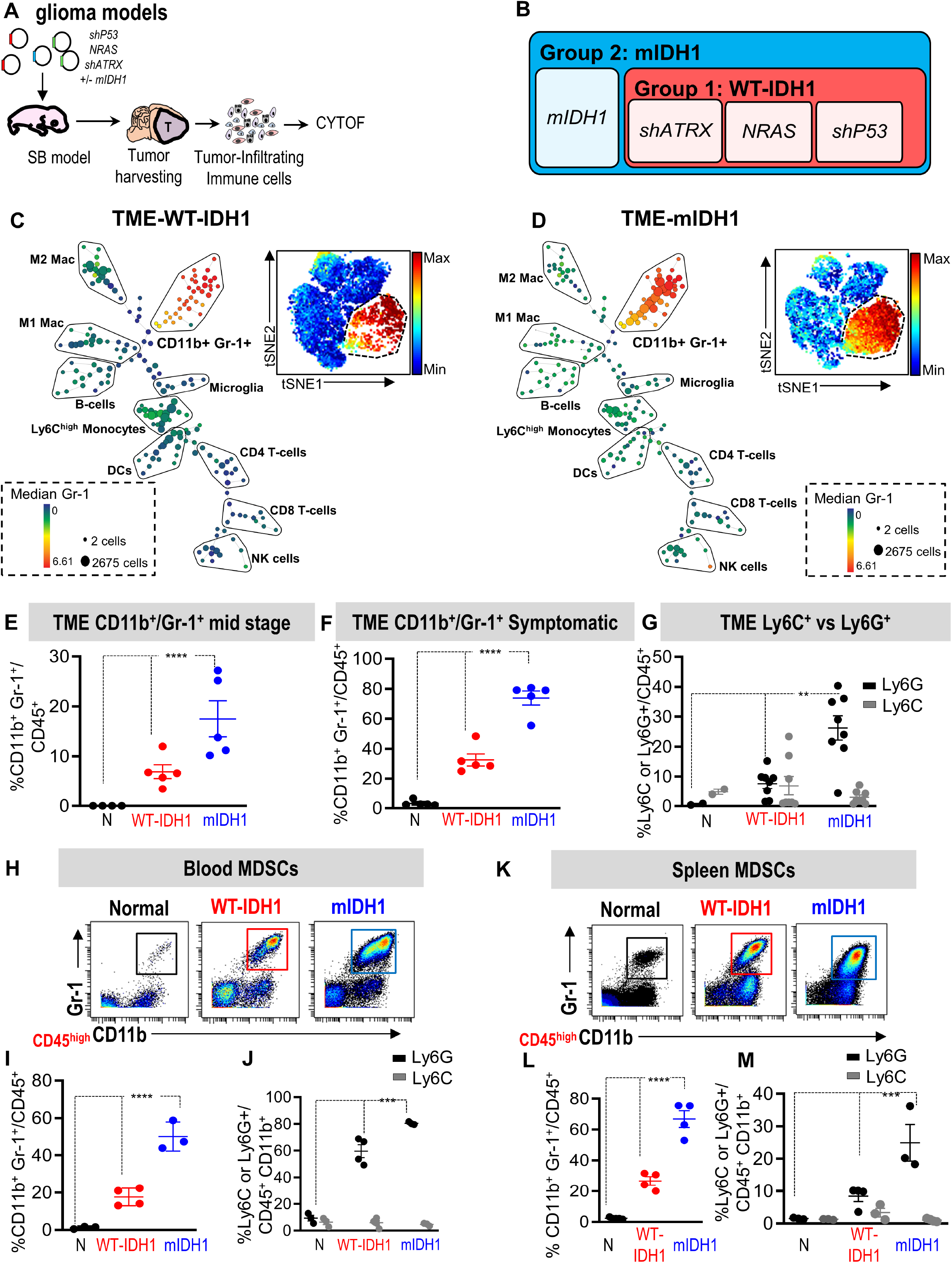
*mIDH1* glioma model has the highest expansion of granulocytic myeloid cells (CD45^high^/CD11b^+^/Ly6G^+^). **(A)** Experimental design of sleeping beauty (SB)-based tumor development, immune cells isolation and CyTOF analysis. **(B)** Schematic representation illustrating the two groups of neurospheres generated with different genetic lesions using the SB model. **(C, D)** SPADE analysis of mass cytometry (CyTOF) data represents immune cell composition within the TME of SB induced *wtIDH1* **(C)** or *mIDH1* **(D)** tumors. Insets are viSNE visualizations of high-dimensional mass cytometry data; color intensity represents the Gr-1 expression level. **(E, F)** After neurospheres implantation, myeloid-derived suppressor cells were identified as CD45^high^/CD11b^+^/Gr-1^+^ cells within the TME at mid-stage of tumor implantation and at symptomatic stage. The percentage of CD45^high^/CD11b^+^/Gr-1^+^ cells was the highest in *mIDH1* tumors. **(G)** Phenotypic characterization of CD45^high^/CD11b^+^/Gr-1^+^ population into granulocytic or monocytic based on the expression of Ly6G or Ly6C at symptomatic stage. The majority of the CD45^high^/CD11b^+^/Gr-1^+^ in the *mIDH1* glioma TME are CD45^high^/CD11b^+^/Ly6G^+^ (granulocytic) cells. **(H, I)** CyTOF analysis of CD45^high^/CD11b^+^/Gr-1^+^ cells in blood from mice with no tumor (normal), and SB induced *wtIDH1* or *mIDH1* glioma. **(J)** Phenotypic characterization of circulating CD45^high^/CD11b^+^/Gr-1^+^ cells according to expression of Ly6C or Ly6G. **(K, L)** CyTOF analysis of CD45^high^/CD11b^+^/Gr-1^+^ cells from the spleens of mice with no tumor, and SB induced *wtIDH1* or *mIDH1* glioma. **(M)** Phenotypic characterization of splenic CD45^high^/CD11b^+^/Gr-1^+^ cells according to expression of Ly6C or Ly6G. N: normal, \**P<0.05, ** P <0.01, *** P <0.005, **** P<0.0001*. Analysis of variance (ANOVA).

### *mIDH1* glioma development results in profound remodeling of BM hematopoiesis

The expansion of the granulocytic myeloid cells within the TME, blood, and spleen suggested that *mIDH1* tumor growth induces profound alterations in the bone marrow (BM) hematopoiesis. Therefore, we examined the changes in hematopoiesis within the BM and spleen of the SB-induced murine glioma model. The absolute number and relative proportion of hematopoietic stem cells (HSCs) were decreased in the BM from *mIDH1* tumor-bearing mice (2.2-fold, < 0.05, Fig. 2A, B). In contrast, BM from *mIDH1* tumors contained higher numbers and frequencies of myeloid progenitors (MPs) (%MP ~35%, < 0.05, Fig. 2C). In the spleen, there were no differences in the frequency of HSCs between the *wtIDH1* and *mIDH1* tumor-bearing animals (Supplementary Fig. S5A, B). However, spleens from *mIDH1* tumor-bearing animals showed more than two-fold increase in MPs compared to *wtIDH1* tumors (*P<0.05*, Supplementary Fig. S5A, C). There was also a higher frequency (2-fold) of granulocyte-macrophage precursors (GMPs) in both the BM and spleen of *mIDH1* tumor-bearing animals (*P*<0.05, Fig. 2D, Supplementary Fig. S5D, F). Despite the increase in MPs and GMPs, there was no significant change in the frequency of common lymphocyte progenitors (CLPs) between the two groups (Fig. 2E, Supplementary Fig. S5E, G). The increased production of granulocytes, MPs, and GMPs in BM and spleen supports the activation of the granulocytic myeloid differentiation pathway in the *mIDH1* tumor-bearing animals (Fig. 2F).

**Figure 2.**
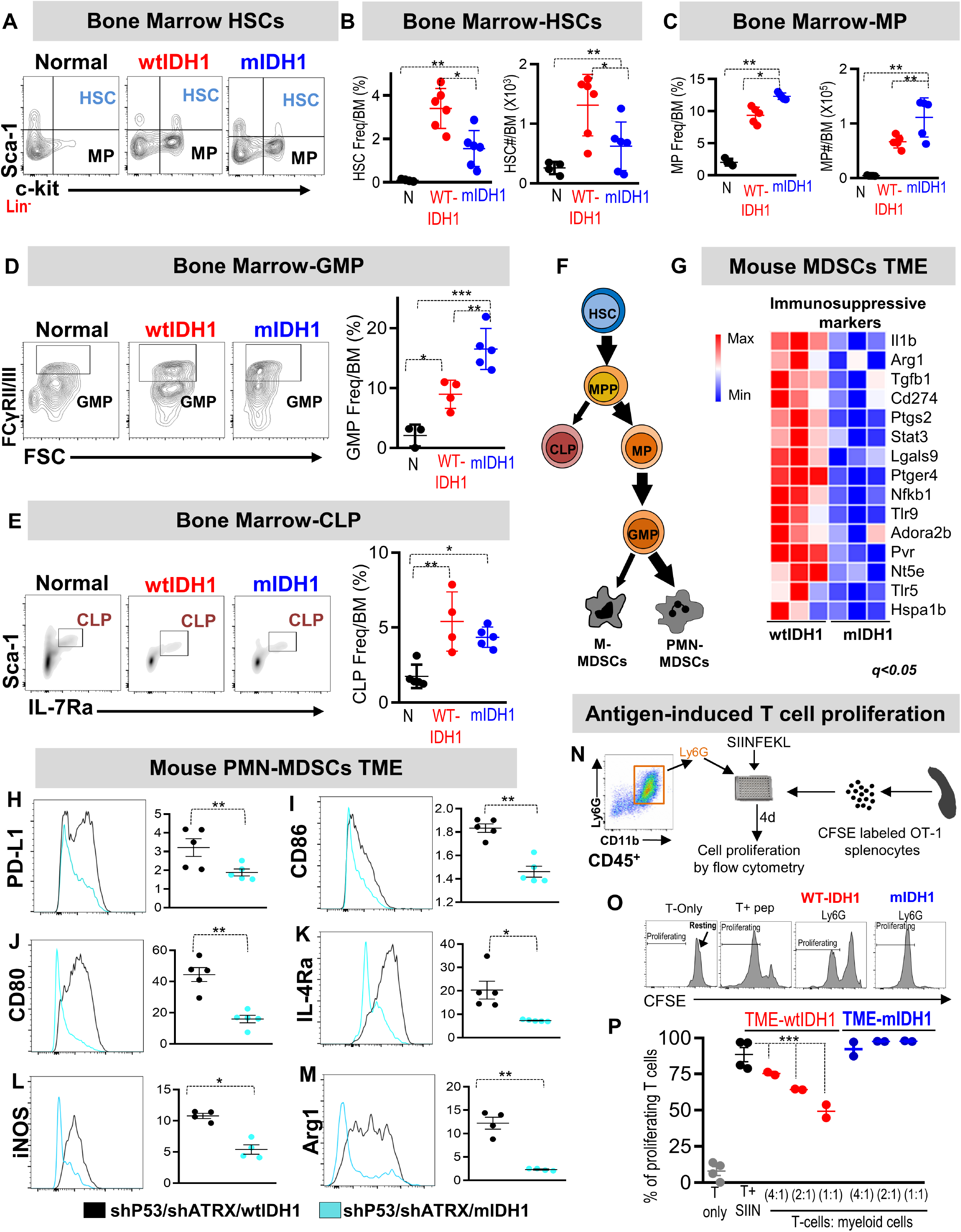
Phenotypic, molecular and functional characterization of myeloid cells lineages in *mIDH1* glioma. **(A)** Representative flow cytometry plots showing the percentage of hematopoietic stem cells (HSCs) (Lin^-^/c-Kit^+^/Sca-1^+^) and myeloid progenitors (MP) (Lin^-^/c-Kit^-^ /Sca-1^+^) in BM from normal mice, and mice implanted with *wtIDH1* or *mIDH1* neurospheres. **(B, C)** Quantification of the number and frequency of **(B)** HSCs and **(C)** MPs from the BM of naïve mouse, and tumor-bearing animals at symptomatic stage. Mice with *mIDH1* tumor have a lower number and frequency of HSCs but higher frequency of MPs. Flow cytometry analysis of the frequency of **(D)** Granulocyte-macrophage progenitors (GMPs) (Lin^-^/IL-7Rα^-^/c-kit^+^/Sca-1^-^ /CD34^+^/FcyRII/III^high^) and **(E)** Common lymphocyte progenitors (CLPs) (c-kit^low^/Sca-1^low^/Lin^-^ /IL-7Rα^+^) in the BM from naïve and *wtIDH1* or *mIDH1* tumor-bearing animals. *mIDH1* tumor had a higher frequency of GMPs but not CLPs in the BM compared to the *wtIDH1* group. **(F)** Schematic diagram representing the shift in hematopoiesis towards granulocytic-lineage development in the bone marrow of *mIDH1* tumor-bearing mice. Thick arrows represent predominant developmental pathways in *mIDH1* tumor bearing mice. **(G)** Heat map showing the normalized expression of genes related to PMN-MDSCs immunosuppressive signature in granulocytes from *wtIDH1* or *mIDH1* TME. CD45^high^/CD11b^+^/Ly6G^+^ cells from *mIDH1* TME have lower expression of immune suppressive markers. **(H-M)** Flow cytometry analysis of immunosuppressive/co-stimulatory markers in the CD45^high^/CD11b^+^/Ly6G^+^ population within TME of *wtIDH1* (black) or *mIDH1* (blue) TME. Granulocytes from *mIDH1* tumors have lower expression of all tested markers. **(N)** Schematic of the *in vitro* T-cell proliferation assay to analyze immune suppressive properties of CD45^high^/CD11b^+^/Ly6G^+^ cells. CD45^high^/CD11b^+^/Ly6G^+^ cells were co-cultured with CFSE-labeled splenocytes from *Rag2/OT-1* transgenic mouse. Cultures were stimulated with 100 nM SIINFEKL peptide for 4 days, after which proliferation was analyzed by flow cytometry **(O)** Representative flow plots showing CFSE staining of unstimulated splenocytes (T only), splenocytes undergoing rapid proliferation in response to SIINFEKL (T+ SIIN), and the effect of SIINFEKL-induced T-cell proliferation in the presence of CD45^high^/CD11b^+^/Ly6G^+^ cells from the TME of *wtIDH1* or *mIDH1* tumors. **(P)** Flow analysis of the inhibitory potential of CD45^high^/CD11b^+^/Ly6G^+^ cells from TME of *wtIDH1* or *mIDH1* tumor. CD45^high^/CD11b^+^/Ly6G^+^ cells from TME-*wtIDH1* tumors are immunosuppressive, whereas CD45^high^/CD11b^+^/Ly6G^+^ cells from TME of *mIDH1* tumors did not suppress T-cell proliferation. \**P<0.05*, \*\**P<0.01*, \*\*\**P<0.005*, One-way ANOVA.

### Molecular and functional characterization of *mIDH1* tumor-infiltrating granulocytes

A hallmark of the glioma microenvironment is the accumulation of immunosuppressive immune cells, which in part is elicited by MDSCs infiltration in the tumor (25). Thus, we investigated the molecular and functional characteristics of the expanded CD45^high^/CD11b^+^/Ly6G^+^ in *mIDH1* tumors. First, we examined the surface expression of T-cell suppressive/co-stimulatory molecules commonly expressed by PMN-MDSCs in *wtIDH1*, and *mIDH1* tumors. Interestingly, in the CD45^high^/CD11b^+^/Ly6G^+^ cells from *mIDH1* TME, the expression level of immunosuppressive as well as costimulatory molecules, such as PD-L1, CD86, CD80, IL-4Rα, iNOS, and Arginase1, was lower compared to the Ly6G^+^ cells from *wtIDH1* TME (Fig. 2 H-M). In order to characterize the molecular differences between mIDH1 and wtIDH1 tumorinfiltrating CD45^high^/CD11b^+^/Gr-1^+^, we purified these cells from tumor-bearing animals of these two groups, and performed transcriptome analysis. Granulocytes from *mIDH1* tumors have decreased expression of all immunosuppressive genes associated with the PMN-MDSCs signature, such as *Il1β*, *Arg1*, *Tgfb1*, *CD274*, and *Stat3* (Fig. 2G). Genes which mediate innate immunity, such as *tlr2*, *tlr7*, and *tlr8*, as well as antigen presentation and processing, were upregulated in granulocytes from *mIDH1* TME (Supplementary Fig. S6A). To further investigate the functions of the differentially expressed genes, we performed gene set enrichment analysis (GSEA) (56). The most significantly enriched gene sets in the *mIDH1* group were related to the immune response (Supplementary Fig. 6B). *Tlr7*, *tlr8*, and *tlr13* were upregulated, whereas genes important for monocyte differentiation (*cebp-β*, and *Irf8*), as well as immunosuppressive genes in myeloid cells (*Il1β*, and *Arg1*), were downregulated (Supplementary Fig. 6B). Collectively, these results suggest that the expanded granulocytes in *mIDH1* tumor exhibit downregulation of immunosuppressive PMN-MDSCs gene signature.

Immunosuppressive PMN-MDSCs are characterized by their ability to suppress T cells’ expansion, therefore we examined the influence of tumor-infiltrating CD45^high^/CD11b^+^/ Ly6G^+^ myeloid cells on antigen-specific T-cell proliferation. We isolated granulocytes by flow cytometry from gliomas expressing *wtIDH1* or *mIDH1* and co-cultured them with carboxyfluorescein diacetate succinimidyl ester (CFSE)-labeled splenocytes from OT-1 mice (Fig. 2N). Isolated granulocytes were next stimulated with the cognate ova albumin peptide SIINFEKL (SIIN) to induce OT-1 splenocyte proliferation (Fig. 2N) (57). In response to SIIN, T-cells proliferation spiked to ~90 ± 4.9% in the control condition (T+ SIIN) (Fig. 2 O, P). When CD45^high^/CD11b^+^/Ly6G^+^ myeloid cells from the TME of *wtIDH1* tumors were added to the culture at different ratios (1:1, 2:1, 4:1), the percentage of proliferating T-cells was reduced to ~48%, 64%, and 75%, respectively (*P<0.001*; Fig. 2O, P). Interestingly, CD45^high^/CD11b^+^/Ly6G^+^ cells isolated from the TME of *mIDH1* gliomas did not suppress antigen-specific T-cell proliferation at any co-culturing ratio (Fig. 2O, P). Together, these results suggest that the granulocytes from the *wtIDH1* tumor are inhibitory and thus can be defined as PMN-MDSCs, whereas the expanded granulocytes in *mIDH1* TME are not inhibitory and cannot be classified as PMN-MDSCs.

To further investigate the impact of *mIDH1* in the context of *ATRX* and *TP53* inactivating mutations on the phenotype and function of CD45^high^/CD11b^+^/Ly6G^+^ cells, we performed an *ex-vivo* MDSC differentiation assay. We isolated BM cells from WT animals and co-cultured them with *wtIDH1*, or *mIDH1* NS (2:1; BM cells: neurospheres; respectively) (Supplementary Fig. S7A). After seven days, we analyzed the frequency of granulocytes formed in each condition. Results showed that in the *mIDH1* NS co-culture, there was a higher frequency of CD45^+^/CD11b^+^/Ly6G^+^ cells compared to those generated by co-cultures with *wtIDH1* NS (63.3 ± 2.1% vs. 85.3 ± 1.5% for wtIDH1 vs. mIDH1, respectively, p <0.05 Supplementary Fig. S7B). Granulocytes in the *mIDH1* NS co-culture expressed lower levels of Arg1, PD-L1 and CD80 compared to those from *wtIDH1* NS co-culture, (Supplementary Fig. S7C-E). The immunosuppressive property of the *ex vivo*-generated CD45^+^/CD11b^+^/Ly6G^+^ cells was investigated by *in vitro* suppression assays. Consistent with the *in vivo* experiments, CD45^+^/CD11b^+^/Ly6G^+^ cells generated by co-culturing with *wtIDH1* NS (1:1) inhibit T-cell proliferation, whereas those generated from *mIDH1* NS were not inhibitory (Supplementary Fig. S7F, G). Collectively, these data validate that a tumor-derived factor is responsible for developing the non-immunosuppressive granulocytes in *mIDH1* glioma TME.

### Single-cell transcriptomes analysis reveals neutrophils and pre-neutrophils are the major granulocytic lineages in *mIDH1* TME

Single-cell RNA-seq provides a powerful approach to dissect heterogeneous immune cells, by providing a transcriptome-wide profiling at the cellular level (58). We used this approach coupled with mass cytometry to compare the molecular differences of the TME-derived granulocytes on an individual cell basis (Fig. 3A). Two biological replicates of purified immune cells from TME of *mIDH1* mice (8,881 cells) and TME of *wtIDH1* mice (7,408 cells) were sequenced at an average depth of ~20,000 reads per cell. After filtering cells with low overall expression and/or high mitochondrial gene expression, we performed dimensionality reduction and unsupervised cell clustering using methods implemented in the Seurat software suite independently for *mIDH1* and *wtIDH1* datasets to identify distinct cell populations (51). In *mIDH1*, we identified 14 distinct cell clusters expressing known markers of major immune cell types (Fig. 3B). Granulocytes formed the largest population and consisted of three clusters, which we refer to as C1, C2, and C3 (Fig. 3B; Supplementary Fig. S8B, D). C1 displayed high expression of PMN-MDSCs-related genes such as *Il1*b and *Arg2*, two major immunosuppressive factors previously used to define PMN-MDSCs in cancer models (59,60) (Fig. 3C; Supplementary Fig. S8B, D). C2 constitutes the largest granulocytic cluster, expressing high levels of *S100a8*, inflammatory cytokines involved in the recruitment of CD8^+^ T-cells (*Ccl3*, and *Ccl4*), and cell cycle genes such as (*Cdc20*, and *G0s2*) (Fig. 3B; Supplementary Fig. S8B, D) (61). C3 expressed genes involved in neutrophil maturation (*Cebpe* and *Mpo*; Fig. 3C, Supplementary Fig. S8B, D). These results suggested that C1 represents the true PMN-MDSCs population within the TME of *mIDH1* tumors, whereas C2 and C3 correspond to pre-neutrophils and neutrophils, respectively. Interestingly, in *wtIDH1*, there was only one cluster of granulocytes (C7; Supplementary Fig. S8A, C). This cluster expressed PMN-MDSC related genes such as *Il1*b and *Arg2*, similar to cluster C1 in TME-*mIDH1* (Supplementary Fig. S8A, C). To confirm that granulocytes in *wtIDH1* tumor (C7) are similar to C1 granulocytes in the *mIDH1* tumor, we integrated the *wtIDH1* and *mIDH1* tumor-infiltrating immune cell datasets using functionality available in the Seurat3 package (50). Jointly clustering *wtIDH1* and *mIDH1* tumor-infiltrating immune cells showed that the granulocytic population formed three clusters corresponding to the C1, C2, and C3 clusters found in the TME of *mIDH1* tumors (Supplementary Fig. S9A). Interestingly, the majority of granulocytes from the TME of the *wtIDH1* tumor clustered with C1 cluster of granulocytes from *mIDH1* tumors (Supplementary Fig. S9B, C), suggesting that these cells share a common origin and have a similar function.

**Figure 3.**
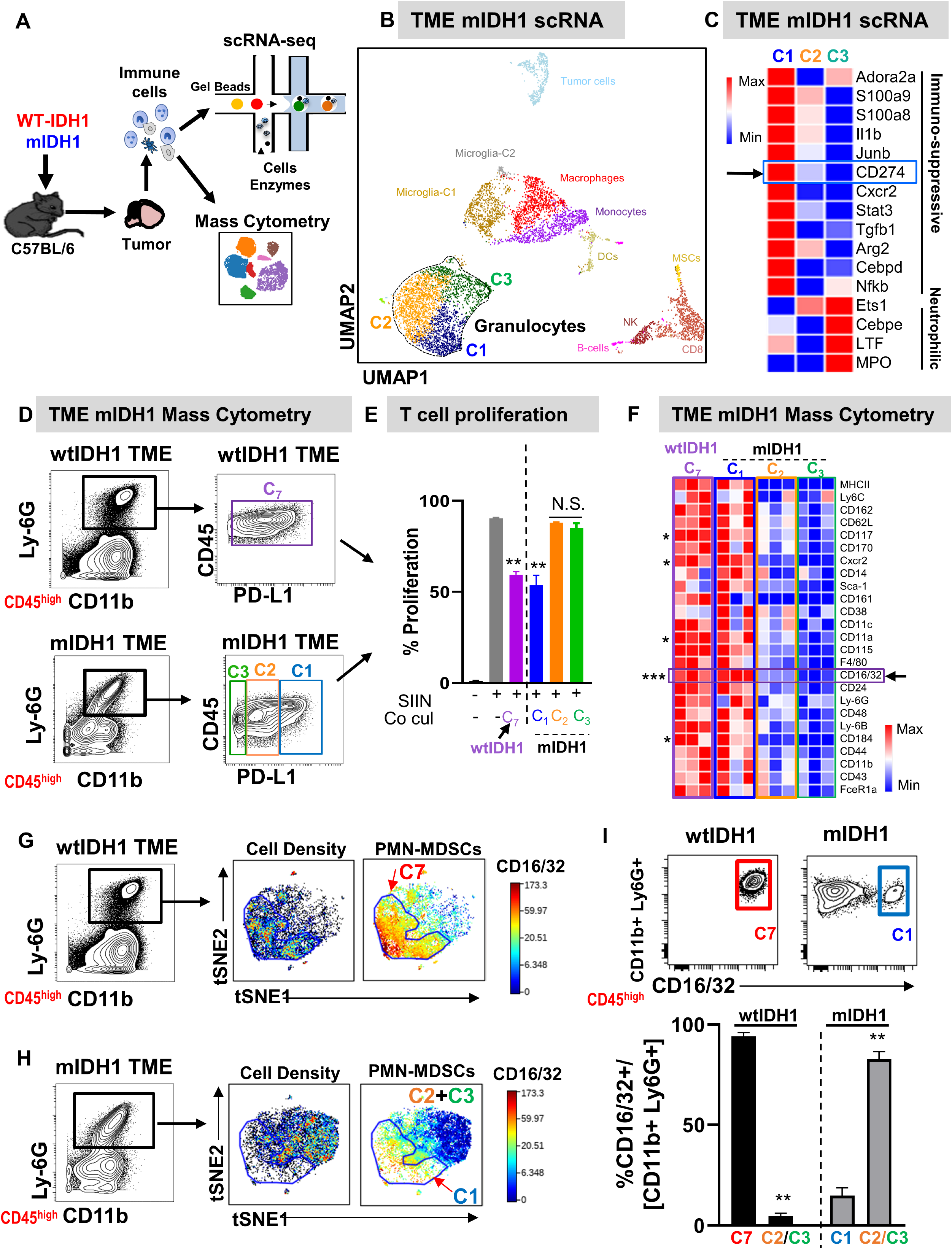
Neutrophils and pre-neutrophils are the major granulocytes population in *mIDH1* tumors. **(A)** Schematic overview of single-cell-seq and mass cytometry (CyTOF) analysis of immune cells purified from TME of SB *wtIDH1* and *mIDH1* tumor-bearing animals. **(B)** Combined Seurat analysis of immune cells from *mIDH1* shown in UMAP projection results in various distinct clusters of TME immune cells. Granulocytes were identified in three clusters in the *mIDH1* TME (C1, C2, C3) (N=2) **(C)** Heatmap of differentially expressed immunosuppressive and neutrophil-related genes between the three granulocytic clusters of *mIDH1* TME group. C1 has a high expression of immune suppressive genes, whereas C2 and C3 have lower expression. C3 has a high expression of characteristic neutrophil genes such as *MPO*, *CEBPE*, and *ETS1*. Arrow reperesent *CD274* expression. **(D)** Mass cytometry analysis of granulocytes from TME of *wtIDH1* (Top panel) or *mIDH1* (bottom panel) tumors. Based on PD-L1 expression, granulocytes from *wtIDH1* tumor clustered in one cell population whereas in *mIDH1*, PD-L1 (CD274) expression, separated the granulocytes into three clusters corresponding to clusters C1, C2, and C3 of the scRNA-seq data. **(E)** T-cell proliferation analysis to assess the inhibitory potential of myeloid cells cluster (C7) from *wtIDH1* tumors or C1, C2, and C3 myeloid cells clusters isolated from *mIDH1* tumors. C7 myeloid cells cluster from the TME of *wtIDH1* tumors and C1 myeloid cells cluster from *mIDH1* tumor TME are immunosuppressive, whereas C2 and C3 from *mIDH1* TME did not suppress T-cell proliferation. **(F)** Heat map representing CyTOF normalized expression level of myeloid biomarkers within granulocytes from the TME of *wtIDH1* tumors or C1, C2, and C3 clusters from the TME of *mIDH1*. CD16/32 (arrow) is reproducibly expressed in immunosuppressive PMN-MDSCs. **(G, H)** Unsupervised clustering and viSNE visualization of granulocytes from *wtIDH1* **(G)** or *mIDH1* **(H)** tumors using biomarker shown in **(F)**. Circumscribed area indicates PMN-MDSCs from *wtIDH1*, which expressed a high level of CD16/32. **(I)** Proportion of PMN-MDSCs (% of total granulocytes) from TME of *wtIDH1* or *mIDH1* TME. The majority of granulocytes from *wtIDH1* (~90%) correspond to PMN-MDSCs, whereas only 12-16% of granulocytes from *mIDH1* belong to PMN-MDSCs. \**P<0.05*, \*\**P<0.01*. Student’s *t*-test

To further determine if C1 represents the true PMN-MDSCs within TME-*mIDH1*, we looked for a cell surface marker differentially expressed in C1 that would allow us to isolate cells from this cluster and assess their function. Among the top 50 differentially expressed genes in cluster C1, *CD274* (PD-L1) was the only cell surface marker that showed high expression in C1 compared to moderate and low expression in C2 and C3, respectively (Fig. 3C; Supplementary Table S4). Based on *CD274* expression, we reclassified the granulocytic populations of both TMEs. We found that there was one granulocyte population in the *wtIDH1* tumor and three distinct granulocytic populations in the *mIDH1* tumor, corresponding to C1, C2, and C3 (Fig. 3D). We FACS sorted each population based on the expression of PD-L1 and determined their inhibitory potential by performing a T-cell proliferation assay. Similar to the results obtained with granulocytes from *wtIDH1* TME (C7 cluster), co-culturing with C1 cells from *mIDH1* glioma results in ~40% inhibition of T-cell proliferation (p < 0.05), whereas C2 and C3 from *mIDH1* TME did not inhibit T-cell proliferation (Fig. 3E). These results validate the inhibitory potential of C1 as the true PMN-MDSCs cluster in *mIDH1* glioma TME.

### CD16/32 is a specific marker that defines immunosuppressive PMN-MDSCs in mice

Our next goal was to identify a specific myeloid biomarker that would allow us to distinguish the inhibitory PMN-MDSCs from the non-inhibitory granulocytes. To identify a unique marker, we used a myeloid-specific antibody panel (Supplementary Table S1) that measures ~30 myeloid-specific parameters simultaneously and used it to perform CYTOF analysis of the granulocytic clusters found in *mIDH1* and *wtIDH1* glioma TME. We found that granulocytes from *wtIDH1* and the C1 cluster from *mIDH1* TME have similar expression of the majority of the myeloid biomarkers, with the FC-gamma receptor family (CD16/32) upregulated in all inhibitory PMN-MDSCs (Fig. 3F). CD16/32 is expressed on macrophages and granulocytes, and its expression is correlated with maturation and function (62,63). To confirm the correlation between PMN-MDSCs phenotype and CD16/32 expression, we performed unsupervised clustering and dimensionality reduction which showed that granulocytes from *mIDH1* TME clustered differently than granulocytes from the *wtIDH1* TME (Fig. 3G, H). We observed high CD16/32 expression exclusively in all granulocytes from *wtIDH1* and only in C1 from *mIDH1* tumors (Fig. 3G, H). The majority of granulocytes from *wtIDH1* TME (~90%) express high levels of CD16/32, while only ~12% of granulocytes form *mIDH1* have high expression of CD16/32, the majority of which belongs to cluster C1 (p < 0.05, Fig. 3G-I). This suggests that CD16/32 is a unique myeloid marker for immune-suppressive PMN-MDSCs in mice.

### PMN-MDSC gene signature is expressed in a higher proportion of tumor-infiltrating immune cells in human *wtIDH1* glioma

We then performed scRNA-seq of immune cells isolated from primary tumor samples of patients with *wtIDH1* and *mIDH1* glioma. We collected 18 tumor samples from a total of 7 patients (8 samples from *wtIDH1* tumors, 10 samples from *mIDH1* tumors). After performing immune cell purification and applying quality controls, a total of 9,765 cells from *wtIDH1* tumors and 17,452 from *mIDH1* tumors were analyzed. All patients’ clinical data are listed in Supplementary Table S5. *wtIDH1* and *mIDH1* datasets were analyzed separately using Seurat to reproduce cell type labels (Fig. 4A-D). Myeloid clusters were the most abundant immune cells infiltrating both the *wtIDH1* and *mIDH1* human gliomas (Fig. 4A-D). In *wtIDH1* tumors, there was a total of 5 clusters of myeloid cells (Myeloid C1-C5) (Fig. 4A, C); all of which expressed the granulocyte colony-stimulating factor receptor, a receptor that labels cells from granulocytic lineages (*CSF3R*) (Supp Fig. S9D). Only one cluster (C4) expressed high microglial genes (*C1Qc*, *AIF1*) (Fig. 4A, C). In the *mIDH1* tumor, there were 4 clusters expressing *CSF3R* (Supp Fig. S9E).

**Figure 4.**
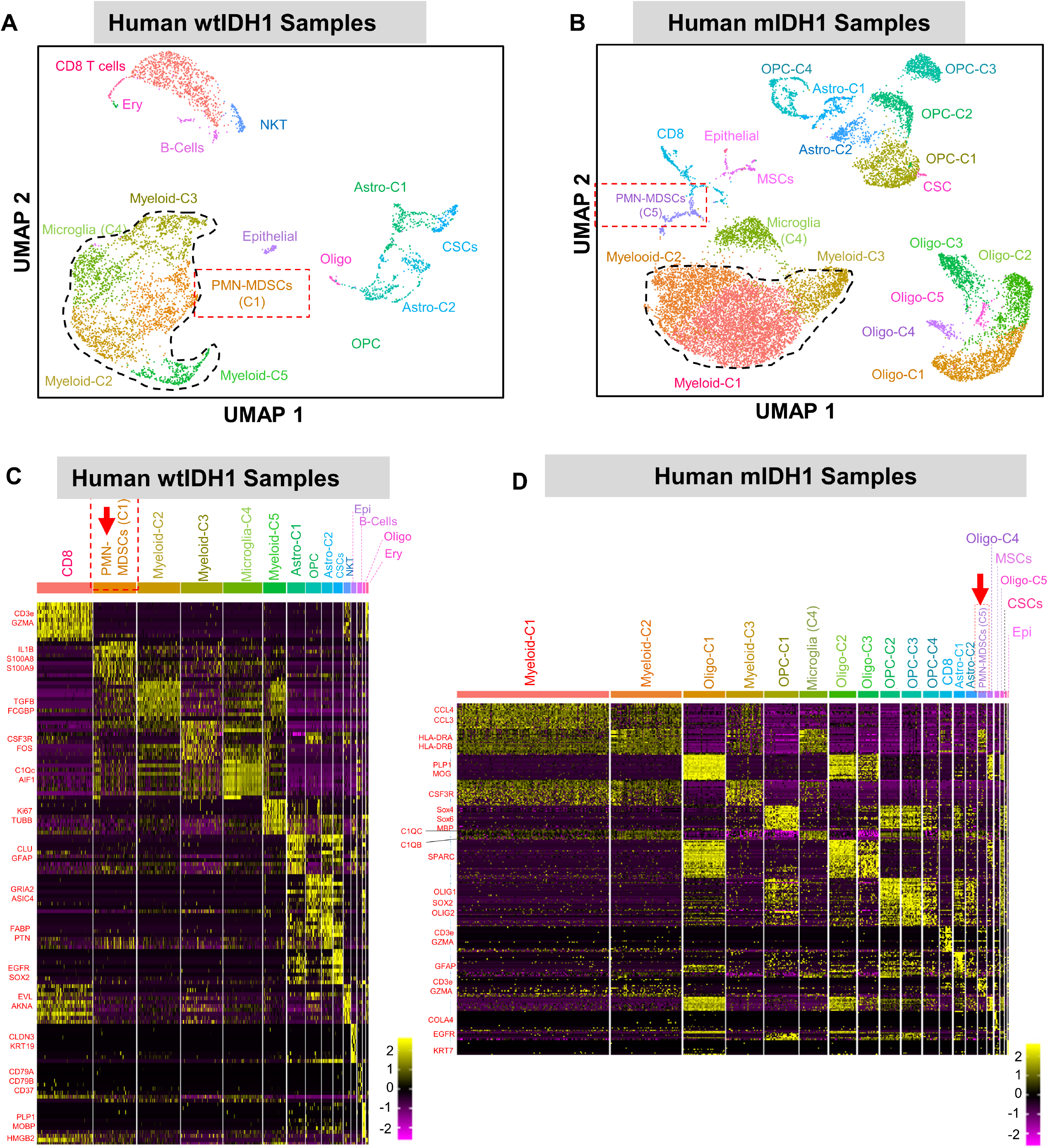
Single-cell RNA-seq analysis of primary human gliomas expressing *wtIDH1* or *mIDH1*. **(A, B)** Seurat analysis of immune cells from **(A)** *wtIDH1* or **(B)** *mIDH1* primary tumor samples shown in UMAP projection results in various distinct clusters. In total, there were five myeloid clusters in *wtIDH1* tumors. The PMN-MDSCs cluster formed the major cell population in *wtIDH1* but not in the *mIDH1* group. **(C)** Heat map showing the differentially expressed genes in each cluster of *wtIDH1* tumor samples. **(D)** Heat map showing the differentially expressed genes in each cluster of the *mIDH1* tumor samples.

Similar to the mouse data, the largest cluster (Myeloid-C1) expressed *CCL3*, and *CCL4* (Fig. 4B, D). Myeloid cluster C2 (Myeloid-C2) expressed antigen presentation-like genes (*HLA-A* and *HLA-DRA*), whereas Myeloid-C3 expressed granulocytes differentiation genes (*CSF3R*, *NEAT1*; Fig. 4D). C4 cluster express the microglia-like genes (*C1QB*, *C1QC*). We then investigated the expression of the PMN-MDSCs gene signature (*IL1β*, *S100a8*, *S100a9, ARG1, TGFβ1*) within each cluster in *wtIDH1* and *mIDH1* myeloid cells, separately (Fig. 5A, B). In *wtIDH1* tumors, Myeloid-C1 expressed high score for the PMN-MDSCs gene signature, whereas in the *mIDH1* group, myeloid-C5 expressed the highest score of PMN-MDSCs signature (Fig. 5A, B, Supp Fig. S9D, E). Nearly 17% of all immune cells in *wtIDH1* tumors were PMN-MDSCs, whereas in the *mIDH1* glioma TME, PMN-MDSCs were only 2.5% of the total immune cells (Fig. 5C). These results are consistent with our preclinical model data which showed that PMN-MDSCs are present at higher fraction in the TME of *wtIDH1* glioma than in the TME of *mIDH1* glioma.

**Figure 5.**
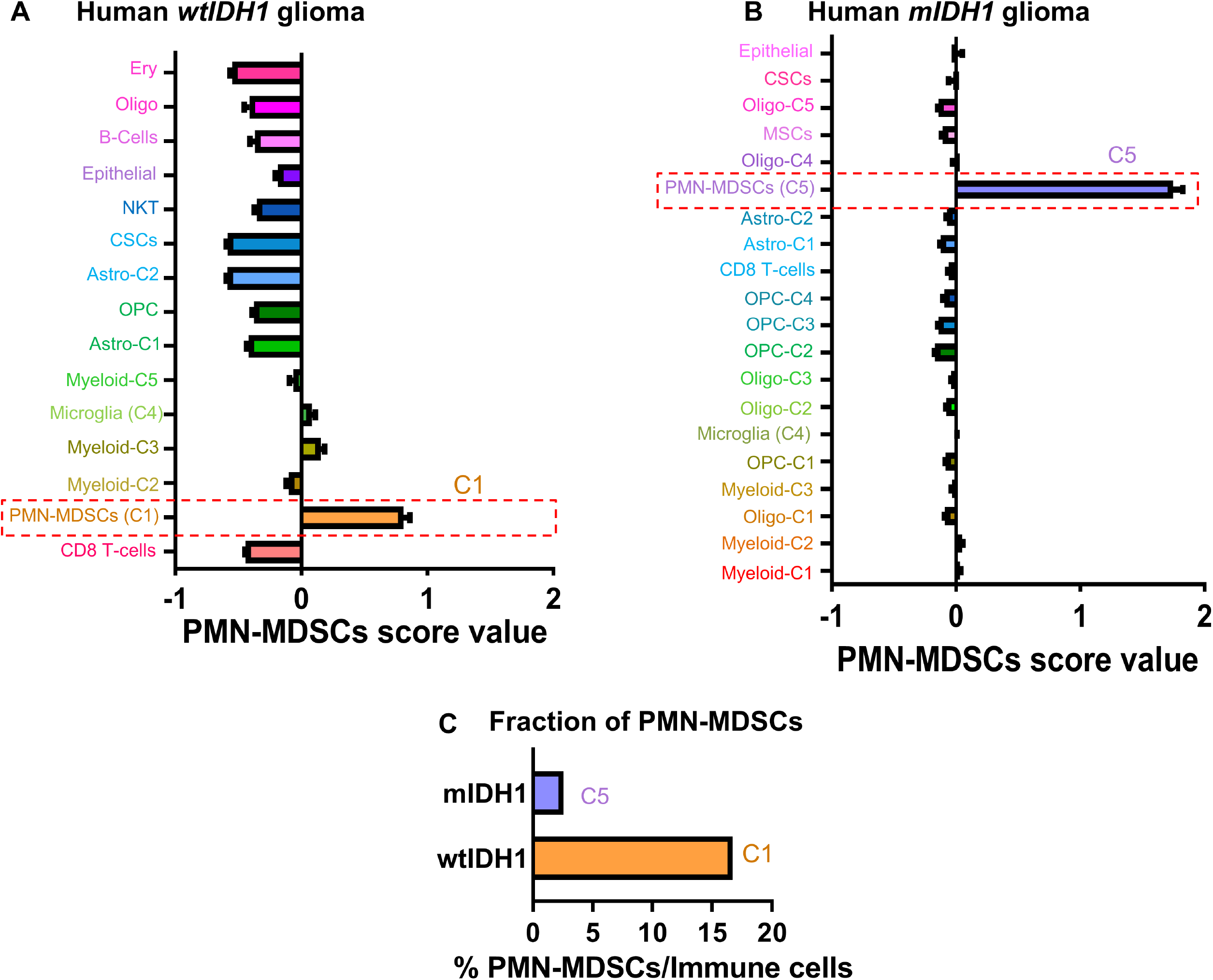
mIDH1 gliomas exhibit lower frequencies of tumor infiltrating PMN-MDSCs. **(A)** Bar plots showing the relative PMN-MDSCs score in each cluster of primary human samples from **(A)** *wtIDH1* or **(B)** *mIDH1* tumors. Cluster C1 in the *wtIDH1* and cluster C5 in the *mIDH1* exhibit the highest expression of PMN-MDSC gene signature. **(F)** Quantification of the percentage of tumor-infiltrating PMN-MDSCs/total immune cells calculated from scRNA-seq data in *wtIDH1* and *mIDH1* human gliomas.

### G-CSF is the major cytokine that is epigenetically regulated in *mIDH1* glioma

Our data demonstrate that the majority of the granulocytes found in *mIDH1* TME are non-inhibitory granulocytes. We hypothesized that the change in granulocytes’ phenotype and function is due to epigenetic reprogramming which affects cytokines’ expression in the *mIDH1* glioma cells. Therefore, we profiled cytokines known to influence myeloid differentiation in conditioned media (CM) collected from *wtIDH1*, or *mIDH1* cultured mouse glioma neurospheres. The expression of the majority of the cytokines was downregulated in CM from the *mIDH1* group compared to CM from *wtIDH1* (Fig. 6A; Supplementary Fig. S10). GM-CSF, CXCL1, CXCL10, IL-5, MIP-2, IL-6, and TNF-α were among the downregulated cytokines in *mIDH1* CM (Fig. 6A; Supplementary Fig. S10). However, G-CSF, RNATES (CCL5), IL-33, and SCF were the only cytokines that were upregulated in *mIDH1* CM (Supplementary Fig. S10). To investigate if these cytokines were epigenetically regulated by *mIDH1*, we used a ChIP-seq dataset that we have recently reported (NCBI’s Omnibus identifier: GSE99806), which was obtained using SB-generated neurospheres from *mIDH1* and *wtIDH1* glioma (64). Gene promoters enriched for H3K4me3 peaks are generally associated with transcriptional activation. Among the genes encoding the cytokines that were upregulated in CM from *mIDH1*, only *CSF3* showed a dramatic change in the peak enrichment for H3K4me3 mark around the promoter region (Fig. 6B). Moreover, quantitative ELISA confirmed that the level of G-CSF was increased in *mIDH1* compared to *wtIDH1* in both the mouse serum of tumor-bearing animals, as well as in CM from cultured mouse and human neurospheres (~5-fold, P <0.05, Fig. 6C-E). In addition, TCGA-LGG analysis revealed that *CSF3* gene expression level was significantly upregulate in *mIDH1* glioma patients harboring *ATRX* and *TP53* mutation as compare to wtIDH1-LGG patients (Figure 6F). The increased level of G-CSF in *mIDH1* could explain the significant expansion of granulocytes in *mIDH1* tumor-bearing mice as well as the enrichment of the cytokine signaling pathway triggered by G-CSF receptor expressed on myeloid cells.

**Figure 6.**
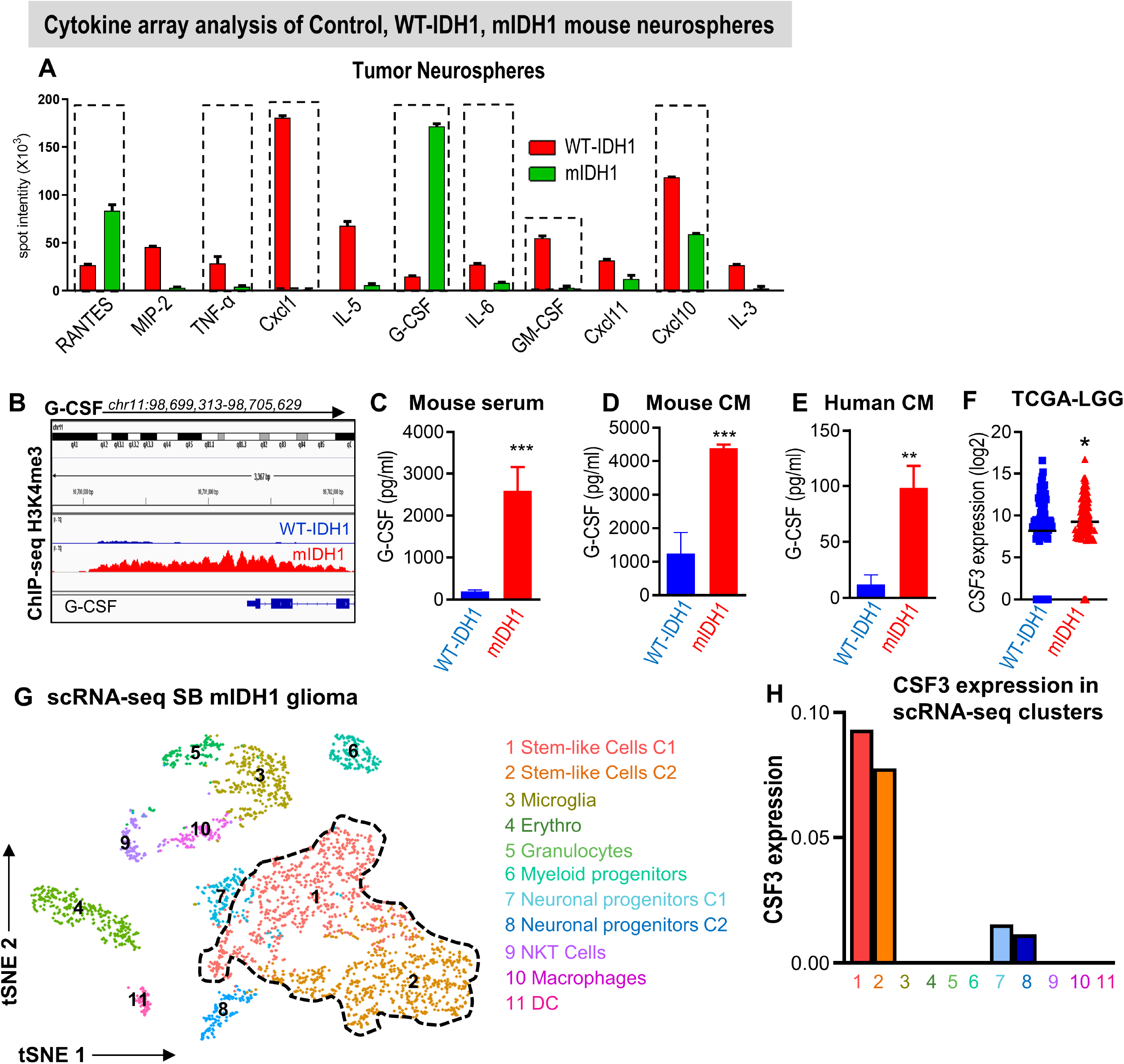
G-CSF is the major epigenetically regulated cytokine expressed by glioma stemlike cells. **(A)** Cytokine array analysis performed on conditioned media from *wtIDH1*, or *mIDH1* neurospheres. Bar graphs show the pixel intensity of each spot. Dash boxes indicate the candidate cytokines that may contribute to the myeloid phenotypic change. **(B)** H3K4me3 occupancy at specific genomic regions of *CSF3* gene. Reads were visualized using IGV, and differential peaks (FDR < 0.05) in *mIDH1* neurospheres are represented in red compared to *wtIDH1* neurospheres in blue. **(C-E)** Quantitative ELISA of the G-CSF level in mouse serum of tumor-bearing animals **(C),** conditioned media from mouse neurospheres cultures **(D),** and conditioned media from human neurospheres cultures expressing *wtIDH1* or *mIDH1* **(E). (F)** Analysis of *CSF3* (G-CSF) gene expression for *mIDH1* glioma patients harboring *TP53* and *ATRX* mutation (n=99), and *wtIDH1* (n=82). RNA-seq data was obtained from TCGA (Xena browser platform). Graph displays the log2 expression value of *CSF3* mRNA. **(G, H)** Combined Seurat analysis shown in tSNE projection of whole tumor cells from *mIDH1* GEMM of glioma results in various distinct clusters of tumor cells. The expression of *CSF3* was analyzed between the clusters. Stem-like cells were the major clusters that have the highest *CSF3* expression. (N=2) * *P<0.05, \**\* *P<0.01, \**\*\* *P<0.005*

To uncover the source of G-CSF within gliomas, we performed scRNA-seq of whole tumors (~5000 cells) isolated from SB induced *mIDH1* glioma model. Results show that the majority of G-CSF was expressed by the major tumor cell clusters, which belong to stem-like cells and express genes such as *sox2*, *sox4*, *tcf4* (Fig. 6G, H). These results indicate that the undifferentiated tumor cells established by the developing *mIDH1* tumor express high levels of G-CSF, resulting in the phenotypic remodeling of the granulocytic population to non-immunosuppressive neutrophils.

### G-CSF neutralization restores the immunosuppressive properties of CD45^high^/CD11b^+^/Ly6G^+^ in *mIDH1* TME

If the CD45^high^/CD11b^+^/Ly6G^+^ phenotype in *mIDH1* is dependent on G-CSF expression, blocking G-CSF should reverse their immune-permissive phenotype. We depleted G-CSF in the *mIDH1* tumor-bearing animals using the αG-CSF neutralizing antibody (Fig. 7A). Treatment with αG-CSF significantly reduced the serum levels of G-CSF in *mIDH1* tumor-bearing mice (~86% and ~92% decrease in day 14 dpi and 21 dpi, respectively; p < 0.01) and decreased the frequency of granulocytes in the TME, spleen, and BM (~40%, ~63%, and ~10%, respectively, p < 0.05 Fig. 7B, C). Interestingly, G-CSF neutralization remodeled the CD45^high^/CD11b^+^/Ly6G^+^ compartment within the TME of *mIDH1* tumor-bearing animals to an inhibitory phenotype, which results in ~35% inhibition of T-cell proliferation (Fig. 7D). Moreover, G-CSF neutralization shortened the MS of *mIDH1* (but not the *wtIDH1*) tumor-bearing animals (MS = 30dpi vs. MS = 18dpi; p <0.05, in control vs. α-GCSF, respectively, Fig. 7E). This demonstrates that the non-inhibitory properties of granulocytes in the *mIDH1* glioma bearing mice is driven by high expression of G-CSF.

**Figure 7.**
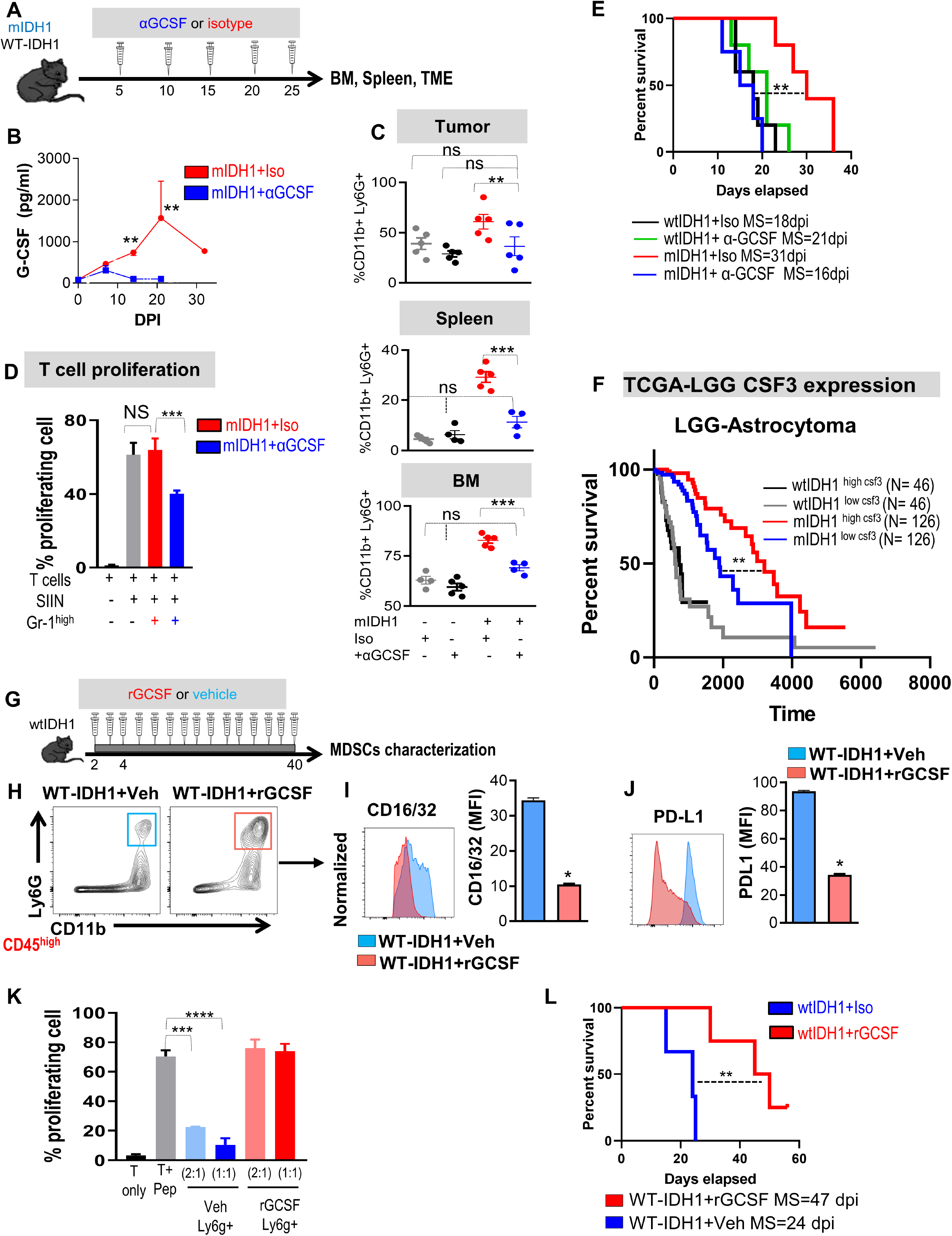
G-CSF neutralization restores the immunosuppressive potential of CD45^high^/CD11b^+^/Ly6G^+^ cells in *mIDH1* glioma and shortened the MS of mDH1 tumorbearing mice. **(A)** Schematic showing experimental design of G-CSF neutralization in *wtIDH1* and *mIDH1* tumor-bearing mice. **(B)** Quantitative ELISA analysis of G-CSF level from serum of *mIDH1* glioma bearing mice treated with either isotype (red) or αG-CSF (blue). **(C)** Flow analysis of CD45^high^/CD11b^+^/Ly6G^+^ cells from tumor, spleen, and BM of tumor-bearing mice treated with either isotype or αG-CSF. G-CSF neutralization decreased the percentage of CD45^high^/CD11b^+^/Ly6G^+^ cells in *mIDH1* tumor-bearing mice but not in the *wtIDH1* tumor. **(D)** T-cell proliferation assay demonstrated that G-CSF neutralization restored the immune suppressive potential of CD45^high^/CD11b^+^/Ly6G^+^ in *mIDH1* tumor-bearing mice. **(E)** Kaplan-Meier survival analysis of mice implanted with either *wtIDH1* or *mIDH1* neurospheres treated with isotype or αG-CSF. Neutralization of G-CSF decreased the MS of *mIDH1* tumor-bearing mice, but it did not affect the MS of *wtIDH1*tumor bearing mice. **(F)** Kaplan-Meier survival analysis of TCGA-LGG astrocytoma *wtIDH1* or *mIDH1* patients with high vs low level of *CSF3* expression. **(G)** Experimental design of recombinant G-CSF (rG-CSF) or vehicle administration in *wtIDH1* tumor-bearing animals. **(H)** Flow analysis of granulocytes from the tumor of *wtIDH1* tumor-bearing animals treated with vehicle or rG-CSF. **(I, J)** Flow analysis of CD16/32 and PD-L1 expression on granulocytes isolated from *wtIDH1* tumor-bearing animals treated with vehicle (blue) or rG-CSF (Red). **(K)** Flow analysis of the inhibitory potential of CD45^high^/CD11b^+^/Ly6G^+^ cells isolated from TME of *wtIDH1* tumor-bearing mice treated with vehicle (blue) or with rG-CSF (red). CD45^high^/CD11b^+^/Ly6G^+^ cells from TME of *wtIDH1*+ veh tumors are immunosuppressive, whereas CD45^high^/CD11b^+^/Ly6G^+^ cells from TME of *wtIDH1*+rG-CSF tumors did not suppress T-cell proliferation. **(L)** Kaplan-Meier survival analysis of animals bearing *wtIDH1* tumors treated with either isotype (blue) or rG-CSF (red). rG-CSF treatment increased the MS of *wtIDH1* tumor-bearing animals (*wtIDH1*+ iso MS=24 dpi, *wtIDH1*+rGCSF MS=47 dpi). * *P<0.05*, ** *P<0.01*, *** *P<0.005*. ANOVA

### *CSF3* expression is associated with favorable outcome only in LGG patients harboring mIDH1

Tumor-derived G-CSF has been shown to promote tumor progression and enhanced metastasis in several tumor types (65,66). To evaluate the effect of G-CSF in LGG patients, we turned to the TCGA data and analyzed the median survival (MS) of all tumor types available in the context of G-CSF (*CSF3*) expression (i.e. MS of patients expressing high vs low levels of *CSF3*). We used the UCSC-Xena browser to identify the differences in survival between patients with high or low levels of *CSF3* expression. Interestingly, we found that LGG was the only tumor type in which a significant difference existed between the MS of patients with high vs low *CSF3* expression (Supplementary Fig. S11). Moreover, when we stratified the astrocytoma patients according to *mIDH1* and *CSF3* expression, we found that enhanced median survival related to high *CSF3* expression was solely observed in patients with *mIDH1* tumors (Fig. 7F). There were no significant differences in MS between high vs low expression of other cytokines responsible for PMN-MDSCs development such as *CSF1, IL4, IL1β, TNFα*, *IL6* in *mIDH1* patients (data not shown). These results suggest the unique correlation between G-CSF expression in *mIDH1* LGG patients and its impact on patients’ survival.

### Glioma derived G-CSF induces the expansion of neutrophils and prolongs survival in mice

To uncover if G-CSF alone is sufficient to induce the expansion of non-immunosuppressive neutrophils, we dosed *wtIDH1* tumor-bearing mice with recombinant G-CSF (rG-CSF) (Fig. 7G). We used 2 μg/day of rG-CSF to achieve steady-state serum G-CSF concentrations comparable to levels observed in *mIDH1* tumor-bearing mice (Fig. 7B), and similar to reported doses (67,68). Compared to vehicle-treated animals, *wtIDH1* tumor-bearing animals treated with rG-CSF showed expansion of granulocytes in the TME (Fig. 7H). These granulocytes expressed lower levels of CD16/32 and PD-L1 compared to granulocytes from vehicle-treated control (Fig. 7I, J). Interestingly, rG-CSF administration remodeled granulocytes to non-inhibitory phenotype and significantly prolonged the MS of tumor-bearing animals (MS = 47dpi vs. MS = 24dpi; p <0.05, in rGCSF vs. control, respectively, Fig. 7K, L). These results supported the hypothesis that G-CSF leads to the expansion of non-immune suppressive neutrophils and confirmed the association between the increased G-CSF secretion and enhanced prognosis in glioma. To validate the source of G-CSF, *wtIDH1* neurospheres were stably transduced with lentivirus encoding G-CSF (lenti-G-CSF) or with an empty lentivirus vector (lenti-vector). Sucessfully ransduced neurospheres were cultured *in vitro* for 14 days before they were used for implantation (Supp Fig S12). The enhanced G-CSF expression was confirmed by RT-qPCR as well as by ELISA in CM of cultured neurospheres (Supplementary Fig. S12B, C). As expected, results showed that animals implanted with *wtIDH1*-lenti-G-CSF neurospheres had increased CD45^high^/CD11b^+^/Ly6G^+^ in the tumor, spleen, and BM compared to mice implanted with *wtIDH1*-Lenti-vector neurospheres (Supplementary Fig. S12D-F). Collectively, this data demonstrates the effect of tumor-derived G-CSF on the phenotype and function of granulocytes in *mIDH1* glioma.

## DISCUSSION

A salient feature of *mIDH1* gliomas is the epigenetic modifications of the tumor cells’ transcriptome mediated by inhibition of the methylcytosine dioxygenase TET2, and αKG dependent histone demethylases. This results in a genome-wide hypermethylation phenotype and enhanced survival (69,70). Our data shows that *mIDH1* stem-like glioma cells, through epigenetic modification, exhibit enhanced G-CSF expression, a strong inducer of myeloid cell development, proliferation, and mobilization (71–73). Enhanced G-CSF expression triggers a shift in myelopoiesis favoring the production of non-immunosuppressive granulocytes that infiltrate the *mIDH1* glioma TME. Thus, our study provides evidence of a novel mechanism that contributes to the enhanced median survival in *mIDH1* glioma.

Gliomas are characterized by a profound immunosuppressive milieu that is responsible, at least in part, for hindering the efficacy of immunotherapies (22–24). Previous studies have shown that glioblastoma patients have elevated levels of circulating MDSCs; the majority of which (~82%) belong to PMN-MDSCs, and are associated with poor prognosis (29,74,75). We have recently shown that the PMN-MDSCs are the predominant immunosuppressive myeloid cells which upon depletion, significantly enhanced the efficacy of immunotherapy and prolonged median survival in *wtIDH1* glioma-bearing mice (25). These results highlight the immunosuppressive nature of *wtIDH1* glioma infiltrating PMN-MDSCs. The exact phenotypic, molecular, and functional characterization of MDSCs in *mIDH1* glioma has not been revealed yet. An algorithm defining the true immunosuppressive MDSCs based on phenotypic, biochemical, and functional characteristics has been recently proposed (42). In our GEMM, although *mIDH1* tumor-bearing animals have a higher expansion of CD45^high^/CD11b^+^/Ly6G^+^ cells, transcriptome and phenotypic analysis revealed that they lack the characteristic immune-suppressive gene signature of *bona fide* PMN-MDSCs (41,42).

The exact mechanisms that control the switch between the suppressive vs. non-suppressive phenotype of polymorphonuclear myeloid cells remain unknown. One proposed mechanism is that two signals orchestrate the development of immunosuppressive myeloid cells (76). The first signal is responsible for the recruitment and proliferation of myeloid cells, i.e., GM-CSF, SCF, or G-CSF (76,77). The second signal belongs to inflammatory cytokines such as IL-1β, IL-6, TGF-β, or IL-4 that trigger an inflammatory milieu responsible for the development of the immune-suppressive properties of the tumor-infiltrating myeloid cells (23,43,78). In line with this hypothesis, our data show that a cytokine belonging to the first signal (G-CSF) was epigenetically upregulated in *mIDH1* glioma bearing mice, whilst, all inflammatory cytokines responsible for the induction of MDSCs immunosuppression properties we tested were downregulated in *mIDH1* glioma. In addition, we found that G-CSF was expressed at a higher level in CM collected from cultured human glioma cells expressing *mIDH/ATRX and TP53* inactivating mutations, when compared to CM collected from *wtIDH1* glioma cells. Moreover, TCGA analysis revealed that the expression of *CSF3* was higher in astrocytoma patients harboring *mIDH1* along with inactivation mutations in *TP53*, *ATRX* as compared to *wtIDH1*. Overall, our data demonstrated that *mIDH1* gliomas exhibited enhanced expression of *CSF3*.

It has been recently shown that gliomas are highly heterogeneous tumors, having both intertumor and intra-tumor heterogeneity at the cellular and genomic levels (79–84). The mechanism by which genetic heterogeneity influences the tumor immune microenvironment remains elusive (85–88). The use of scRNA-seq technologies has now provided additional granularity which enables to chart subtypes of glioma immune cellular infiltrates within the glioma microenvironment. This technology enabled us to identify the developmental hierarchies within the glioma infiltrating myelopoietic cell clusters, uncovering novel drivers, and revealing the infiltrating-immune surveillance/evasion cell types relevant to tumors with specific genetic lesions (84,89–91). We thus performed scRNA-seq of the tumor-infiltrating CD45^+^ immune cells harvested from our glioma GEMMs; both *wtIDH1* and *mIDH1*. Transcriptome profiling at the single cell level uncovered the global landscape of glioma-infiltrating myeloid cells and captured their heterogeneity. We discovered that the *mIDH1* glioma-infiltrating granulocytes are comprised of three distinct cell clusters; C1, C2, and C3. We performed differential gene expression analysis between the three clusters and identified that cluster C1 was enriched in immunosuppressive PMN-MDSCs signature marker genes such as *tgfb*, *Il1b*, and *Arg2*. Cluster C2 expressed *S100a8*, *ccl3*, *ccl4*, and cell cycle genes such as *Cdc20*, *G0s2* indicating proliferating neutrophils (pre-neutrophils). Whereas C3 expressed neutrophils maturation genes such as *cebpe*, and *mpo*. In contrast, in the *wtIDH1* glioma model, the tumor-infiltrating granulocytes are comprised of one single cluster expressing genes similar to granulocyte cluster, C1 in the *mIDH1* group.

Since a hallmark of MDSCs, is their inhibition of antigen specific CD8^+^ T cell proliferation, we next aimed to isolate the three newly identified granulocytic clusters encountered within the *mIDH1* TME for functional characterization. An ideal marker would be expressed on the cell surface, and have different expression levels between the three clusters. Interrogation of the differentially expressed genes between C1, C2, and C3 revealed that *CD274* (PD-L1) was expressed at a high level in PMN-MDSCs (cluster C1); compared to intermediate and low levels in the pre-neutrophils (cluster C2) and neutrophils (cluster C3), respectively (Fig. 3C). This allowed us to isolate each cluster and examine their T-cell inhibitory potential. Using this approach, we found that in the *mIDH1* group, cells in cluster C1 were suppressive whereas the cluster C2 and C3 were non-immunosuppressive, whereas in the *wtIDH1* group, granulocytes expressing PD-L1 formed one single cluster of cells (C7), which inhibited CD8^+^ T-cell proliferation. Finally, we performed in-depth high-dimensional mass cytometry characterization and phenotypic analysis of the three granulocytic clusters found in the *mIDH1* TME. An advantage of this approach is that automated clustering is done on data reduced to two tSNE dimensions, which allows visual verification of cluster boundaries at the single-cell resolution. Combined clustering of granulocytes infiltrating the TME of *mIDH1* or *wtIDH1* GEMMs of glioma revealed that the majority of granulocytes from the *wtIDH1* tumor were allocated along with the C1 cluster of the *mIDH1* group. Cytof analysis also led to the identification of CD16/32 as a unique, and reproducible myeloid marker that can be used to distinguish immunosuppressive PMN-MDSCs from non-immunosuppressive granulocytes.

Consistent with results from our GEMMs of glioma, we found that the majority of immune cells infiltrating human gliomas expressing *mIDH1*, exhibit a characteristic myeloid cell’s gene signatures; all of which express the early granulocytic receptor G-CSFR. Interestingly, the largest myeloid cluster in the *mIDH1* glioma group expressed genes (*CCL3*, *CCL4*) similar to cluster C2 in our *mIDH1* GEMM of glioma. Moreover, using a unique PMN-MDSCs gene signature, our data elucidated that human *mIDH1* gliomas exhibit a lower frequency of tumorinfiltrating *bona fide* PMN-MDSCs when compared to the *wtIDH1* glioma patients. Therefore, the molecular and phenotypic features of *mIDH1* on tumor-infiltrating PMN-MDSCs is largely conserved between our mouse model and human samples.

G-CSF was upregulated in the CM of cultured human and mouse *mIDH1* NS. This increase was also significant in the serum from *mIDH1* tumor-bearing mice. We demonstrated that the increased G-CSF expression in *mIDH1* NS was due to epigenetic reprogramming of the tumor cells’ transcriptome, in which the *CSF3* gene showed an enrichment in the deposition of the H3K4me3 mark around the promoter region. We demonstrated that in our *mIDH1* GEMM of glioma, tumor cells express a higher level of *Sox2* and are less differentiated (46); consistent with earlier reports indicated that *IDH1^R132H^* suppresses cellular differentiation (92,93). scRNA-seq revealed that glioma stem-like cells that express *Sox2* and *Sox4* are the major source of G-CSF expression in *mIDH1* tumors. Therefore, the enhanced production of G-CSF is due to the presence of a higher proportion of less differentiated tumor cells in *mIDH1* glioma. In line with this, studies showed that within the glioma niche, glioma stem-like cells are found to be coenriched with myeloid infiltrating immune cells (94,95).

The impact of G-CSF on tumor biology is still a point of debate. Recombinant G-CSF (Filgastim®) is a commercially available product indicated for chemo-induced neutropenia to reverse myelosuppression in cancer patients and it has been shown to reduce inflammation and reverse cognitive impairment in patients with Alzheimer (96,97). In addition, increased G-CSF secretion has been implicated in the recruitment and infiltration of immunosuppressive MDSCs in breast cancer (65) and cervical cancer (66). In contrast, in mice bearing MCA203 sarcomas, G-CSF induced CD11b^+^/Gr-1^hi^ cells that were non-suppressive (98). It has also been shown that the administration of G-CSF leads to the formation of non-immunosuppressive CD45^+^/CD11b^+^/Gr-1^+^ *in vivo* (98) and *in vitro* (99). Our data in *mIDH1* glioma show that the increased production of G-CSF by *mIDH1* glioma stem-like cells caused the expansion of CD45^high^/CD11b^+^/Ly6G^+^ cells that exhibit a non-immunosuppressive phenotype. We also showed that rG-CSF administration to mice harboring *wtIDH1* glioma reversed the immunosuppressive properties of the PMN-MDSCs, leading to increased median survival, similar to the median survival observed for *mIDH1* glioma bearing mice.

In summary, our data demonstrate that *mIDH1* tumor-derived G-CSF reprograms myelopoiesis in glioma, which promotes the production of non-immunosuppressive granulocytes, a feature that can be harnessed to enhance the immunotherapeutic efficacy in *mIDH1* glioma patients (Supplementary Fig S13).

## ACKNOWLEDGEMENTS

We greatly thank Kait Verbal, and Andrea Gunawan for all the help. We thank the support and academic leadership of Dr. Karin Muraszko, and the administrative and technical support of Angela Collada, and Marta Edwards, respectively. The content is solely the responsibility of the authors.

